# Quantitative analysis shows that repair of Cas9-induced double-strand DNA breaks is slow and error-prone

**DOI:** 10.1101/142802

**Authors:** Eva K. Brinkman, Tao Chen, Marcel de Haas, Hanna A. Holland, Waseem Akhtar, Bas van Steensel

## Abstract

The RNA-guided DNA endonuclease Cas9 is a powerful tool for genome editing. Little is known about the kinetics and fidelity of the double-strand break (DSB) repair process that follows a Cas9 cutting event in living cells. Here, we developed a strategy to measure the kinetics of DSB repair for single loci in human cells. Quantitative modeling of repaired DNA in time series after Cas9 activation reveals a relatively slow repair rate (~6h). Furthermore, the double strand break is predominantly repaired in an error-prone fashion (at least 70%). Both classical and microhomology-mediated end-joining pathways are active and contribute to the repair in a stochastic manner. However, the balance between these two pathways changes over time and can be altered by chemical inhibition of DNAPKcs or additional ionizing radiation. Our strategy is generally applicable to study DSB repair kinetics and fidelity in single loci, and demonstrates that Cas9-induced DSBs are repaired in an unusual manner.

## Introduction

The CRISPR-Cas9 nuclease system is a powerful tool for genome editing due to its efficient targeting of specific sequences in the genome (Sander and Joung, 2014). Cas9 endonuclease is directed by a guide RNA to a specific target site in the genome, where it induces a single double-strand break (DSB) (Cong et al., 2013; Jinek et al., 2013; Mali et al., 2013). The break is subsequently repaired by the cellular DNA repair mechanisms that can introduce mutations in the target sequence (Jasin and Haber, 2016). This application of Cas9 has become widely popular to generate mutant alleles of genes and regulatory elements of interest. Despite the broad application, the process of repair of Cas9-induced DSBs has been only partially characterized. For example, it is not known how long it takes before an individual Cas9-induced DSB is repaired, and how error-prone this process is.

Eukaryotic cells have two main pathways for DSB repair: classical non-homologous end-joining (C-NHEJ) and homologous recombination (HR) (Jasin and Haber, 2016). A large proportion of DSBs is repaired by C-NHEJ, which directly rejoins the two DNA ends. This type of repair is thought to be mostly perfect but may lead to insertions or deletions (indels) at the break site (Lieber, 2010). However, estimates of the frequency at which these indels occur are a matter of debate. The amount of precise rejoining has been suggested to range between 35 - 97% depending on the damaging agent and the end structures of the DSBs (Betermier et al., 2014; Jasin and Haber, 2016; Liang et al., 2016, 2016). In contrast, HR is highly precise because it utilizes a homologous template sequence to restore the DNA sequences around the DSB (Greene, 2016). Apart from these two main pathways, there are alternative repair pathways that are highly mutagenic. One of these is microhomology-mediated end-joining (MMEJ), which uses short sequence homologies near the two ends, leading to characteristic small deletions (McVey and Lee, 2008). Current evidence indicates that multiple pathways can contribute to the repair of Cas9-induced DSBs, but the interplay and the relative contributions are largely unclear (Bothmer et al., 2017; van Overbeek et al., 2016). Moreover, the fidelity of these pathways in the context of Cas9-induced breaks is still largely unclear.

Related to this, it is still unknown how quickly a Cas9-induced break is repaired. Several methods have been developed to measure the rate of rejoining of DNA DSBs in the genome. Direct measurements of DSBs include comet assays (Wang et al., 2013) or pulsed-field gel electrophoresis (DiBiase et al., 2000). Time series of these DSB data and mathematical modeling have yielded estimates of overall repair rates (Cucinotta et al., 2008; Woods and Barnes, 2016). After exposure to ionizing radiation, causing hundreds of DSBs per cell, most damage was repaired within 0.25 – 2 h (Kinner et al., 2008). Similar repair rates were measured using immunofluorescent labeling of DSB markers such as γH2A.X, combined with microscopy, flow cytometry or chromatin immunoprecipitation (Forment et al., 2012; Redon et al., 2009; Seo et al., 2012). Recently, methods were developed to directly detect DSBs genome-wide (Canela et al., 2016; Crosetto et al., 2013; Soong et al., 2015; Zhou et al., 2013), but these have not yet been used to follow the kinetics of repair.

Sequencing-based methods offer new opportunities to directly measure the outcome of repair at single-nucleotide level. For example, sequencing of repaired DNA around Cas9-induced DSBs has revealed that each locus has a distinct spectrum of insertions and deletions (indels) (Brinkman et al., 2014; van Overbeek et al., 2016). This spectrum can be modulated by inhibition of C-NHEJ (van Overbeek et al., 2016), suggesting that multiple pathways can act on one DSB.

In principle, accumulation of mutations over time can be used to infer repair rates. A crude estimate based on indel detection suggested that about 15 hours were necessary to repair the majority of the Cas9-induced lesions (Kim et al., 2014). However, precise quantitative kinetics of actual re-joining of DNA ends after the induction of a DSB at a single defined genomic location are missing. One challenge is that perfectly repaired junctions are indistinguishable from DNA that was never broken, which may lead to systematic errors in the rate constants. In addition, with Cas9 the accumulation of indels is also dependent on the cutting rate. Here, we tackle these problems by a combination of mathematical modeling and highly accurate measurements of indel accumulation. Our modeling is based on the principle that the precise kinetics of indel accumulation is dictated by the rate constants of cutting, perfect repair and mutagenic repair. Conversely, these parameters can be inferred from curves of measured indels over time.

The results indicate that repair after a Cas9 cut is remarkably slow, and that error-free repair of the DNA is almost absent. Furthermore, the repair at a single locus is executed by C-NHEJ or MMEJ in a stochastic way with distinct dynamics. The balance between these two pathways can be altered by chemical inhibition or by additional damage elsewhere in the genome. The observation of high levels of imperfect repair may explain the effectiveness of CRISPR-Cas9 for genome editing.

## Results

### A kinetic model of DSB repair

We approached the process of repair of a Cas9-induced DSB as a simple three-state model (Figure 1A). In this model, the *intact* state is the original unbroken DNA sequence that can be recognized by the Cas9/sgRNA complex. After introduction of a DSB by this complex the DNA enters a reversible *broken* state. This state may be repaired perfectly, after which the DNA is susceptible to another round of cutting. Alternatively, an error-prone repair mechanism may introduce a small insertion or deletion (indel) at the break site. The latter results in an irreversible *indel* state that can no longer be recognized properly by the sgRNA and therefore cannot be cut again by Cas9 (see below for validation of this assumption). Hence, in this model there are three reaction steps: cutting, perfect repair and mutagenic repair, each having a specific rate constant that we refer to as *k_c_, k_p_* and *k_m_,* respectively (Figure 1A).

**Figure 1:**
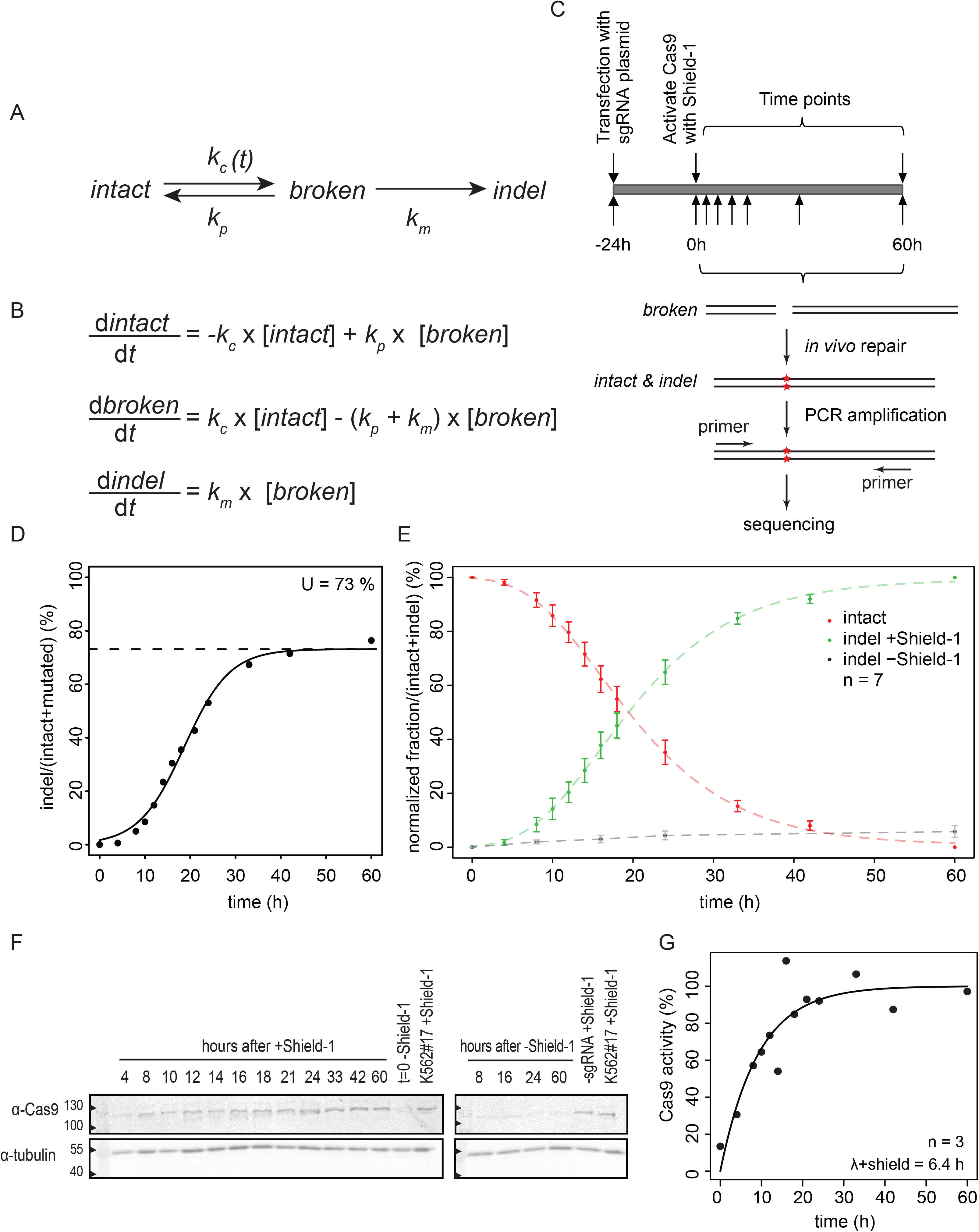
Quantitative analysis of Cas9-induced DSBs. (**A**) Proposed model of DSB repair based on stochastic transitions between intact, broken and indel states. *k_p_* and *k_m_* are rate constants of perfect and mutagenic repair, respectively; *k_c_* is the rate constant of cutting by Cas9. The latter depends on Cas9 activation and is therefore denoted as *k_c_(t).* (**B**) ODEs describing the three reaction steps, with rate constants as parameters. (**C**) Outline of the experimental strategy; see main text. (**D**) Representative time course experiment, showing gradual accumulation of indels. A sigmoid curve was fitted to the data to determine the plateau level at late time points (dashed line), which reflects the transfection efficiency. (**E**). Relative proportions of intact (red) and indel (green) fractions of the LBR2 locus over time. The data points are normalized on to the total indel fraction to correct for the variation in transfection efficiency. Indel fraction in absence of Shield-1 is shown in grey. Average of 7 independent experiments; error bars represent the standard deviation and the dashed lines show fitted sigmoid curves. (**F**) Western blot analysis of Cas9 presence in time. Tubulin was used as loading control. (**G**) The intensities of Cas9 antibody signal were determined by photo densitometry from time points of 3 individual western blots and normalized to a sample incubated for 60 hours with Shield-1 (lanes labeled “K562#17+Shield-1” in (**F**)). ODEs fit was performed to determine the activity score of Cas9 in time.

Our aim was to determine these key descriptors of the repair process for individual Cas9/sgRNA target loci. For this purpose, we captured the model in a set of ordinary differential equations (ODEs) describing the three reaction steps, with the rate constants as parameters (Figure 1B). With this ODE model it is possible to simulate the relative abundance of the three states over time in a pool of cells after activation of Cas9. Such simulations show that activation of Cas9 generally leads to a gradual loss of intact DNA, a transient increase in the broken DNA state, and a gradual increase in the indel state until virtually all DNA is converted to the latter. However, the shape of each curve is determined by the rate constants (Supplementary Figure S1). We reasoned that it should therefore be possible to determine the rate constants by fitting the ODE model to actual time series measurements of one or all three states after Cas9 induction. This approach has successfully been applied to highly similar kinetic models in a variety of research fields (Ay and Arnosti, 2011; Bintu et al., 2016; Cucinotta et al., 2008).

### Quantification of Cas9 cutting and repair rates

We then set out to measure the accumulation of indels in cells in which a specific DSB was introduced by Cas9. To control the timing of DSB formation we established a clonal K562 cell line (K562#17) with a stably integrated construct that encodes a tightly controlled inducible Cas9. To switch Cas9 activity on and off, we fused a ligand-responsive destabilizing domain (Banaszynski et al., 2006) to the Cas9 nuclease. With the small ligand Shield-1, Cas9 can then be reversibly stabilized for transient DSB-induction. K562 cells are capable of activating DNA damage response upon DSBs, although the G1 checkpoint is affected due to a mutated *TP53* gene (Law et al., 1993).

We transiently transfected K562#17 with a plasmid encoding a sgRNA targeting the LBR gene (sgRNA-LBR2). We previously found that this sgRNA effectively induces indels (Brinkman et al., 2014). Twenty-four hours after transfection we stabilized Cas9 by adding Shield-1. Flow cytometry analysis showed that cells 16 h after damage had an ~10% increase in G2 population suggesting a DNA damage response and check point activation (Supplementary Figure S2). We collected cells at various time points, isolated genomic DNA, amplified a ~300-bp region around the sgRNA target site by PCR and subjected the products to high-throughput sequencing to determine the intact and indel fractions (Figure 1C).

The results show a gradual accumulation of indels over time (Figure 1D), indicating that DSBs were introduced and repaired imperfectly. Towards the end of the time course (60 h) the indel frequency reached a plateau of ~70%. This value corresponds approximately to the transfection efficiency, as indicated by the proportion of cells that express GFP after transfection with a GFP-expressing plasmid (Figure 1D, Supplementary Figure S3). We therefore assumed that the plateau value of ~70% represents the total proportion of cells that received the sgRNA and underwent DSB induction and repair. After normalization for this transfection efficiency, the data were highly reproducible over 7 independent replicate experiments (Figure 1E). Cells that were not incubated with Shield-1 showed almost no indels, indicating that the process is dependent on stabilized Cas9 (Figure 1E).

The sigmoid appearance of the measured indel time curves suggested a delayed onset of indel accumulation. This was not observed in any of the initial simulations (cf. Supplementary Figure S1). We reasoned that this discrepancy may be explained by a delayed activation of Cas9 at the beginning of the time series. Indeed, Western blot analysis showed that Cas9 accumulates gradually after addition of Shield-1 (Figure 1F). Assuming that the cutting activity of Cas9 is proportional to its abundance, we modified the computational model to incorporate the gradual increase of Cas9 levels as empirically determined from the Western blot signals (Figure 1G).

### Cutting and repair rate constants in the LBR gene

Next, we fitted the set of ODEs to the measured indel curves for sgRNA-LBR2 (Figure 2A–D). Based on 7 independent replicate experiments, this yielded a cutting rate *k_c_* = 0.11 ± 0.01 h^-1^ for maximum Cas9 expression. This corresponds to a cutting half-life (i.e., the time that would be required to cut 50% of the available target sequences in the absence of repair) of ~5.5 hours. Cutting by Cas9/sgRNA-LBR2 in our system is thus a rather slow process.

**Figure 2:**
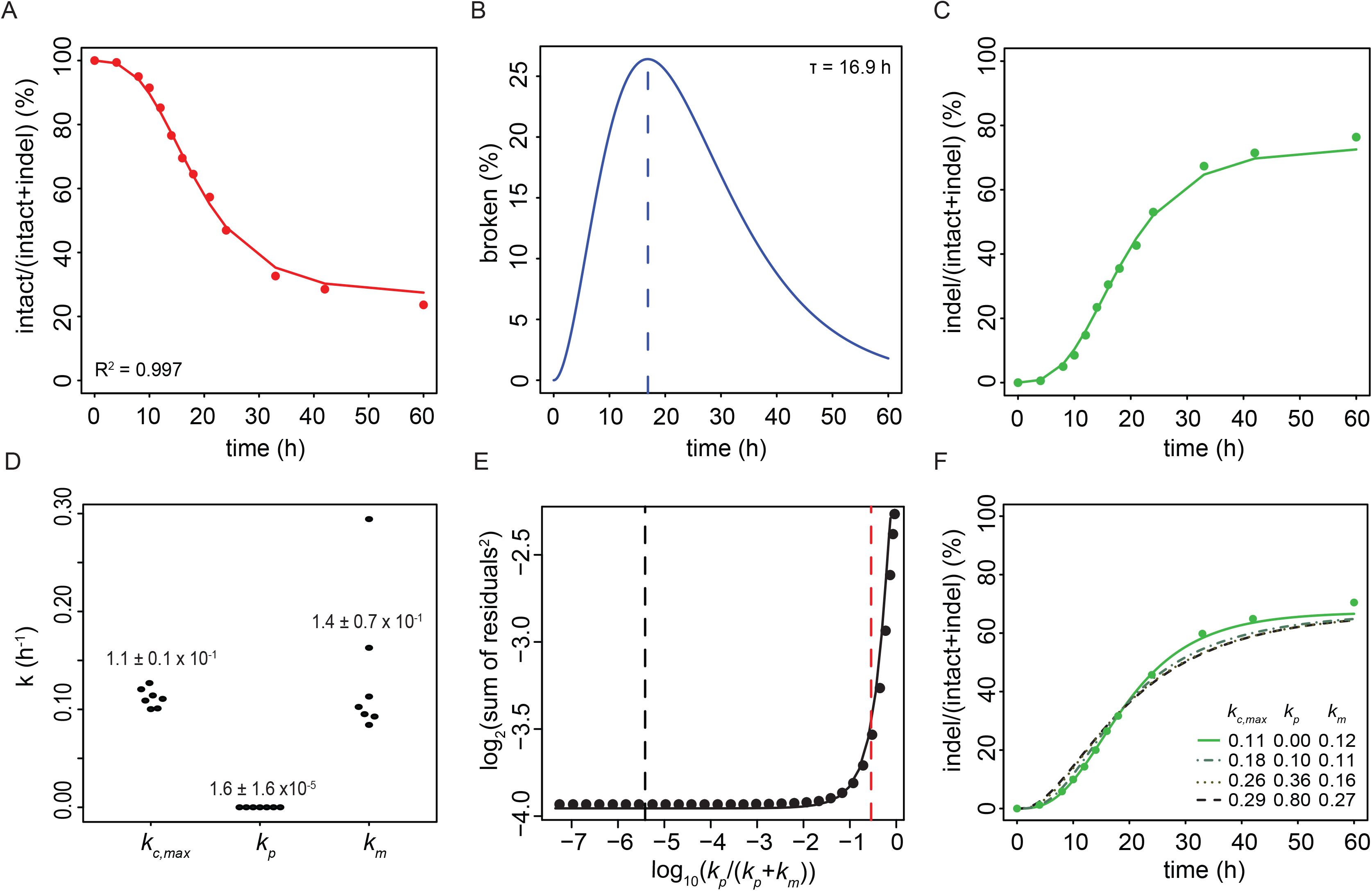
Estimation of cutting and repair rates. Representative time series traces of the intact (**A**), broken (**B**) and indel (**C**) fractions. Measured data (dots) are overlaid with the ODE model fit (solid lines). The percentage of intact and indel traces are relative to the total. The broken fraction is estimated by the model on the basis of the the intact and indel measurements. *t* indicates time of the largest amount of broken DNA. (**D**) Distributions of rate constants from 7 independent experiments (values indicate mean ± SD). (**E**) Estimate of the upper confidence bound of the proportion of perfect repair. Fitting residual errors are shown for various fixed *k_p_/*(*k_p_*+*k_m_*) ratios (7 combined time series). Black dashed line marks the optimal fit; red dashed line marks the *k_p_/*(*k_p_*+*k_m_*) ratio above which the fit becomes significantly worse than the optimal fit (P <0.05, one-sided F-test). This corresponds to *k_p_/*(*k_p_*+*k_m_*) = 0.29. For each *k_p_/*(*k_p_*+*k_m_*) ratio, *k_c,max_* and *k_m_* were adjusted to produce the best fit. (**F**) Examples of model fitting with various combinations of *k_p_* and *k_m_* forced to values that would correspond to models of rapid cycling of cutting and perfect repair (dashed curves). Green dots are measured values, green curve shows unrestrained optimal fit.

Our model fit estimated the rate constant for imperfect repair to be *k_m_* = 0.14 ± 0.07 h^-1^. This corresponds to a half-life of broken DNA of 5.1 hours. Surprisingly, the rate constant for perfect repair was estimated to be *k_p_* = 1.6 ± 1.6 x 10^-5^ h^-1^, which is nearly ten thousand times slower than imperfect repair. This suggests that virtually all repair events at this locus result in the formation of indels, while perfect repair is very rare.

### Robustness of the model

On average the goodness of fit between the model and the measured data was R^2^=0.999. Furthermore, we considered that the parameter estimates were strongly influenced by the modeling of the Cas9 induction. To test this we also modelled a simple step function with various time delays of Cas9 activation. Although the results were quantitatively slightly different (Supplementary Figure S4), the main conclusion remained that repair is slow and error-prone.

While these results point to robustness of the modeling, we were surprised to find the extremely low rate of perfect repair. We therefore conducted further analyses to check whether the model with the low perfect repair rate is indeed the most optimal fit to the data. Specifically, we conducted a “parameter sweep” survey in which we imposed different fixed perfect/mutagenic repair ratios and compared the goodness of fit as measured by the sum of squares of the fitting residuals. This indicated that low *k_p_*/(*k_p_*+*k_m_*) ratios (~10^-6^) indeed yield the best fit. However, the difference with higher ratios were rather minor (Figure 2E–F), and statistical testing revealed that only at *k_p_*/(*k_p_*+*k_m_*) ratios > 0.29 the fit became significantly poorer (P < 0.05, F-test) than at the initially estimated ratio of 10^-6^ We therefore conservatively conclude that the contribution of perfect repair can be at most 29%, although lower ratios are more likely to be correct. The other parameter values (*k_c_*,*k_m_* and the predicted broken fraction) showed only minor variation within this range (Supplementary Figure S5), further attesting to the robustness of the model.

From this parameter sweep analysis follows that the indel accumulation curve is not compatible with a very rapid cycle of cutting and perfect repair with only occasional imperfect repair. In such a scenario *k_p_*/(*k_p_*+*k_m_*) would be close to 1, which yields a poor fit compared to lower values (Figure 2E–F).

### Experimental validation

In addition to these computational tests we validated the underlying assumptions and results of the modeling with several independent biological assays. First, we tested our assumption that the reverse reaction from the indel state to the broken state cannot occur (Figure 1A). For this purpose we isolated several clonal cell lines from K562#17 after transfection with sgRNA-LBR2 and Shield-1 induction of Cas9 for ~20 days. As expected, these clones had acquired one or more indels in the target site and lacked wild-type sequences (Supplementary Figure S6A). We then re-transfected 3 of these cell clones with sgRNA-LBR2 and again activated Cas9. Despite prolonged re-exposure to Cas9/sgRNA-LBR2, we could not detect any change in the indels present in each clone (Supplementary Figure S6B). We conclude that the target site, once it has acquired an indel, is not recognized again by the same sgRNA.

Second, we verified the kinetics of the broken state as predicted by the computational model (which was based on intact and indel frequencies only). The model predicts that the broken fraction peaks at 16.4 ± 1.7 hours, with a maximum of 19.9 ± 6.7% broken DNA (mean values ± SD, n = 7; Figure 2B). To verify this, we established a variant of the ligation-mediated PCR assay for the quantification of DNA breaks at a defined location (Chailleux et al., 2014; Dai et al., 2000; Garrity and Wold, 1992). In this assay, we first denature the DNA and subject it to a primer extension reaction using a primer near the break site. This ensures that all cleavage sites are converted into blunt ends, even if resection of the broken ends has occurred. Next, an adaptor is ligated to the blunted DNA end, followed by PCR with one primer near the break site and a second primer that is complementary to the adaptor sequence (Figure 3A). When analyzed on agarose gels, the samples from cells treated with sgRNA and Shield-1 yielded a band of the expected size (Figure 3B). Analysis of several time series consistently showed that the band intensity increased until 12.7 ± 2.6 h (mean ± SD, n = 3) hours after Cas9 induction, and then decreased again. This is in agreement with the peak time of the broken state as predicted by the model fitting (Figure 3C). Furthermore, the parameter sweep shows that when *k_p_*/(*k_p_*+*k_m_*) approaches 1, the predicted amount of broken DNA at peak time becomes so low (<1%) that we would be unable to measure it (Supplementary Figure S6D-E). Thus, a model consisting of a very rapid cycle of cutting and perfect repair with only occasional imperfect repair is not compatible with our ability to detect broken DNA.

**Figure 3:**
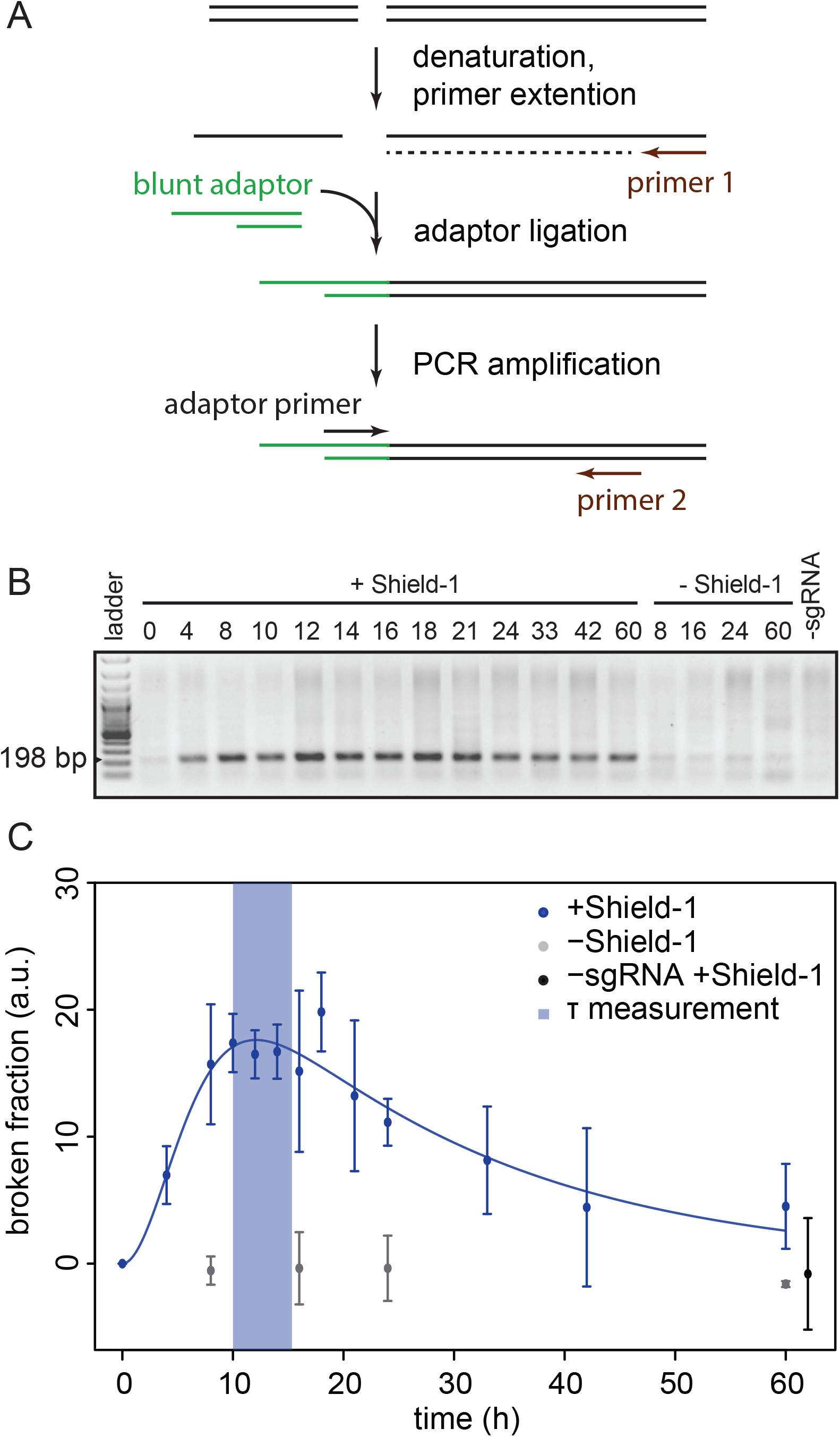
Measurement of broken fraction. (**A**) Schematic view of the LM-PCR assay to detect the broken ends after DSB induction. See text for full explanation. (**B**) Representative agarose gel of the LM-PCR products of a time series. The expected product is 198 bp in size. (**C**) The broken fraction measured as band intensities (data from 7 LM-PCR experiments spanning 3 different time series; values are mean ± SD). Solid blue line shows an ODE curve fit to the LM-PCR data to determine *τ* (blue shading, mean ± SD).

### Two repair pathways active at one locus

Next, we took a closer look at the specific indels that were generated. We and others showed previously that repair of Cas9-induced DSBs produces non-random indel patterns that are specific for the sgRNA (Brinkman et al., 2014; van Overbeek et al., 2016). In our experiments sgRNA-LBR2 yielded predominantly a deletion of 7 bp or an insertion of 1 bp (Figure 4A, B). After 60 hours of Cas9 induction the +1 insertions reached a frequency of 42.3 ± 2.8%, while the −7 deletions accumulated to 13.1 ± 2.0% (Figure 4C). Analysis of 20 clonal lines derived from single cells in which Cas9/sgRNA-LBR2 had been transiently active indicated that the +1 insertion is the predominant mutation but can co-occur with −7 deletion on other alleles in the same cell (Supplementary Figure S6A).

**Figure 4:**
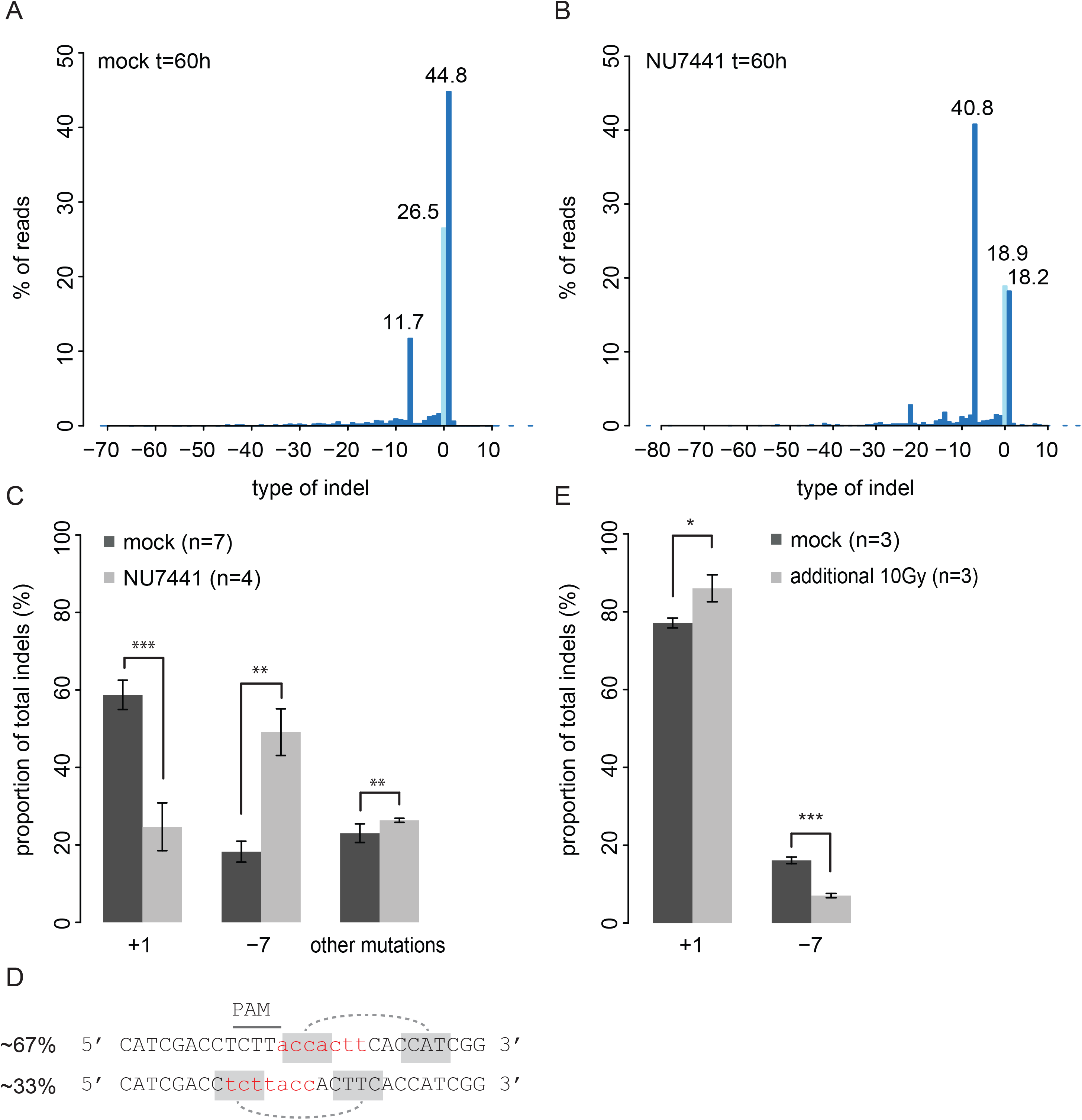
Multiple repair pathways active at one locus. (**A-B**) The spectrum of indels and their frequencies at the LBR2 locus at time point t = 60 h, in cells cultured without (**A**) or with (**B**) 1 μM NU7441. A representative experiment is shown. Light-blue bar: wild-type sequence; dark-blue bars: indels. (**C**) Quantification of indel fraction in the presence (black, n=7) and absence (grey, n=4) of NU7441. All series are normalized to the total indel fraction. Asterisks indicate P-values according to Student’s t-test: * P<0.05, ** P<0.005, *** P<0.0005. (**D**) Percentages represent the proportion of observed −7 sequence reads. Red nucleotides indicate the deleted DNA. Grey shades show possible models for microhomology mediated repair. (**E**) TIDE analysis of +1 insertion and −7 deletion indels after exposure of the cells to 10 Gy of IR at the time of Cas9 induction. Indels fraction is normalized on the total indels

Previous studies have indicated that different indels can be the result of different repair pathways, e.g. C-NHEJ or MMEJ (Hicks et al., 2010; van Overbeek et al., 2016). To explore whether this may be the case for the −7 and +1 indels, we added an inhibitor of DNA-PKcs (NU7441) to the cells. DNA-PKcs plays an essential role in the C-NHEJ pathway but not in MMEJ. We found that in the presence of 1 μM NU7441 the balance of indel type changed: the proportion of the −7 deletion events increased by 3-fold while the +1 insertion diminished by about 2-fold. In addition, more larger deletions arise (Figure 4A, B). It is important to note that the presence of NU7441 inhibitor did not change the cell viability (Supplementary Figure S7A). These results indicate that the +1 insertion is the result of C-NHEJ, while the −7 deletion is not.

MMEJ makes use of microhomologies near the broken ends (Guirouilh-Barbat et al., 2004; Guirouilh-Barbat et al., 2007; Wang et al., 2006). Sequence analysis revealed that the −7 deletion fraction in fact consists of two types of deletions that occur in an approximate ratio of 1:2 (Figure 4D). Both types can be explained by recombination through 3-nucleotide microhomologies, which strongly points to MMEJ as the responsible pathway for the formation of the −7 indel. We conclude that at least two different repair pathways are active and lead to distinct types of mutations at one specific break site.

Interestingly, we found that the ratio between the +1 and −7 indels changes in favor of the +1 insertion when 10 Gy of ionizing radiation (IR) damage was administered just prior to Cas9 induction. In particular, the −7 deletion fraction decreased about 2-fold (Figure 4E). Cell viability did not differ between the control and irradiated sample (Supplementary Figure S7B). The shift in pathway utilization can be either due to additional breaks elsewhere in genome (which may sequester components of the MMEJ pathway), a cell cycle arrest or a combination thereof.

### Delayed activity of MMEJ

Having established that two types of indels are largely the result of separate pathways, we decided to study the kinetics of these pathways in more detail by tracking the +1 and −7 indel frequencies over time. We modified the ODE model by incorporating separate *k_m_* rate constants for each type of indel (Figure 5A). We then fitted this model to the +1 and −7 indel time series data. Inspection of the resulting model revealed a good fit of the +1 curve (mean R^2^ = 0.99), but a poor fit of the −7 curve (mean R^2^ = 0.93; Figure 5C). The deviation of the fitted curve is mostly due to a delay in the −7 indel appearance (Figure 5B). As a consequence, our estimate of the *k_m_* for the −7 indel is less accurate, but we can conclude that the MMEJ pathway exhibits a delayed onset compared to C-NHEJ. Thus, the *k_-7_* rate appears to increase over time, rather than be constant. Possibly the MMEJ pathway is only activated when C-NHEJ fails to be repair a DSB, as has been proposed previously (DiBiase et al., 2000; Guirouilh-Barbat et al., 2004; Guirouilh-Barbat et al., 2007; Wang et al., 2003; Wang et al., 2005).

**Figure 5:**
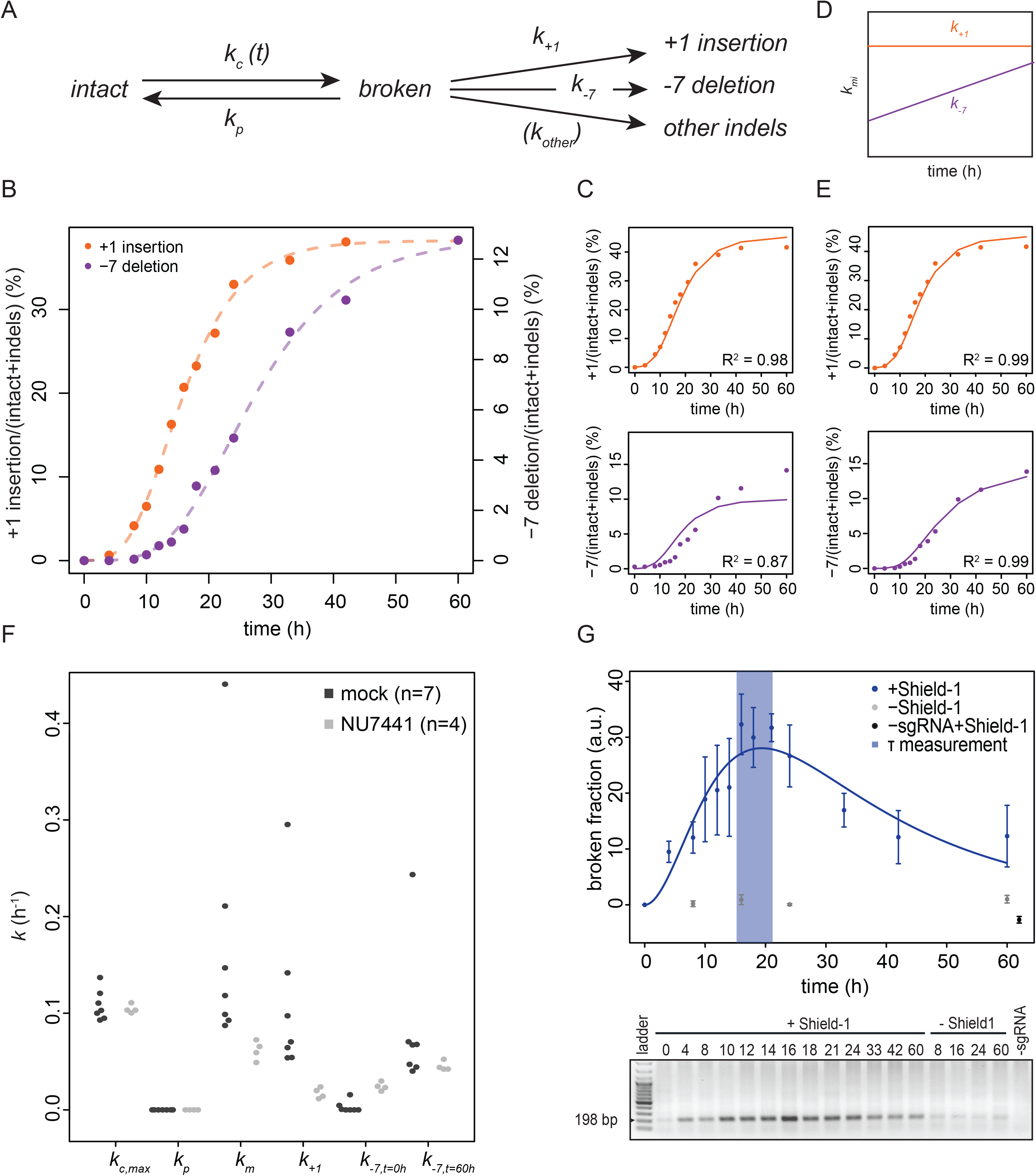
MMEJ has slower repair kinetics than C-NHEJ. (**A**) Kinetic model of Cas9-induced DSB repair assuming that the various indels are generated with a specific repair rate. (**B**) +1 insertion and −7 deletion accumulation in a representative time series. Dashed lines show a Gompertz sigmoid fit. (**C**) Same time series as in (**B**) now overlaid with the multi-indel ODE model fit (solid lines) for the +1 (top) and −7 (bottom) indels. (**D**) Cartoon representation of hypothesized time dependency of the *k+_1_* and *k_-7_* rates added to the fitted model in (**E**). (**E**) Time dependent ODE model fit of the +1 insertion (top) and −7 deletion (bottom) as proposed in (**D**). (**F**) Rate constants without (black) and with (grey) NU7441. The total mutagenic repair rate is represented by *k_m_*; rate constant for the +1 insertion is *k_+1_* and rate constants for the −7 deletion are *k_-7,t=0h_* and *k_- 7,t=60h_* at the start and end of the time series, respectively. *n* indicates the number of time series. (**G**) Broken fraction measurements in presence of NU7441 of 2 independent time series, similar to Figure 3B–C.

We then tested whether the model could be improved by explicitly including such a delayed onset. For simplicity we assumed a linear increase in the production rate of the −7 deletion over time, starting with a rate *k*_-7,*t*=0*h*_ immediately after Cas9 induction and increasing to *k*_-7,*t*=60*h*_ at the end of the time course (Figure 5D). Indeed, a much better fit of the −7 indel curve (mean R^2^ = 0.99) was obtained (Figure 5E). The model fitting estimates *k_-7_* to be nearly zero at the onset of Cas9 induction, while at the end of the time course it is 0.08 ± 0.07h^-1^, which approaches the activity of the +1 indel repair 0.11 ± 0.09 h^-1^ (Figure 5F). Reassuringly, the new model predicted *k_c_, k_p_* and *k_m_* values that are similar to the previous estimates (compare Figure 2D **and** 5F). Taken together, these results strongly suggest gradual activation of MMEJ over time.

### Interplay between C-NHEJ and MMEJ

We wondered whether this gradual increase in MMEJ rate is somehow due to competition with the C-NHEJ pathway. To test this, we performed time course experiments in the presence of NU7441 and repeated the computational modeling. As expected, in the presence of the inhibitor *k_+1_* is reduced dramatically (~8-fold; Figure 5F, Supplementary Table 3), while the cutting rate *k_c_* as well as the perfect repair rate *k_p_* are virtually unaltered. Strikingly, in the presence of NU7441 *k*_-7,*t*=0*h*_ became much higher than in the absence of the inhibitor, while *k*_-7,*t*=60*h*_ remained largely unaffected. Thus, inhibition of DNA-PKcs leads to a more rapid engagement of MMEJ soon after the DSB is introduced.

However, in the presence of NU7441, MMEJ does not fully compensate for the loss of C-NHEJ. The total repair rate is lower, as *k_m_* is reduced and *k_p_* remains close to zero (Figure 5F). A logical prediction is that it takes more time for DSBs to be repaired. Indeed, the model as well as actual measurements show that in the presence of NU7441 the peak time of DSBs is delayed by about 4-7 hours (compare Figure 3B and 5G).

### Cutting and repair rates are similar in three other loci

To investigate whether the rate constants are locus-specific, we performed kinetics experiments for three additional loci (Figure 6A–I; Supplementary Table 4). We designed a second sgRNA (sgRNA-LBR8) targeting a different sequence in the *LBR* gene, 169 bp upstream from sgRNA-LBR2. We also chose a previously reported sgRNA in the *AAVS1* gene (Mali et al., 2013) and additionally designed a sgRNA that targets an intergenic locus. For each target we conducted time series measurements and determined the rate constants by fitting of the ODE model to the total indel fractions.

**Figure 6:**
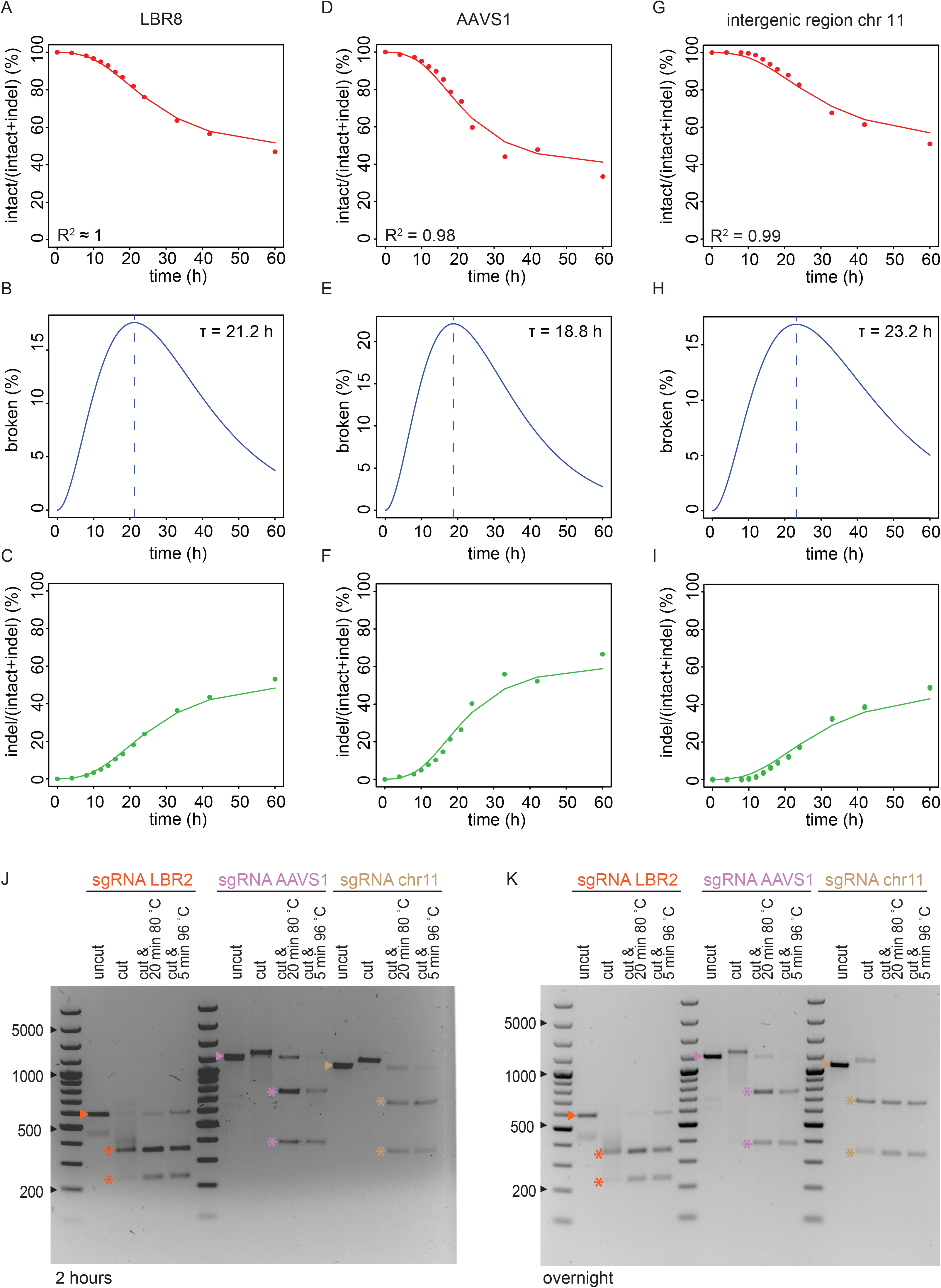
ODE modeling of DSBs at additional loci; Cas9 remains bound to broken ends. (**A-I**) Time series of Cas9 cutting and repair at three other loci (LBR8 (**A-C**), AAVS1 (**D-F**), intergenic region on chromosome 11 (**G-I**)). Relative percentages are plotted of the intact (red) and indel (green) fraction of the measured sequence reads (dots). Solid line indicates the ODE fit of each individual fraction. The broken fraction is inferred from the model and its peak time is indicated by τ. (**J, K**) *In vitro* digestion of DNA fragments by Cas9 and one of the sgRNAs targeting LBR (orange), AAVS1 (purple) or intergenic region on chromosome 11 (brown), respectively. DNA was incubated for either 2 hours (**J**) or overnight (**K**) in the absence (uncut) or presence of Cas9/sgRNA (cut). In some samples Cas9 was subsequently denatured by two different heat treatment protocols as indicated. The expected band for the unbroken DNA is marked by an arrowhead and the expected digestion products are marked by asterisks.

While each target locus shows a different indel spectrum (Supplementary Figure S8), the overall rate constants for the four loci are all in the same range (Supplementary Table 4). The estimated cutting rates (*k_c_*) and mutagenic repair rates (*k_m_*) each vary over a ~1.5-fold range only. We note that for two of the loci the indel frequency did not fully reach a plateau at the end of the time course, which may somewhat compromise the accuracy of the rate constant estimates (see Methods). The minor differences between the four loci may also be due to effects of sequence context, nucleosome positioning (Horlbeck et al., 2016; Isaac et al., 2016) or local chromatin environment. For sgRNA-LBR8 we also measured a time course of the broken state, which corresponded well with the predicted time course according to the ODE model (Supplementary Figure S9).

Strikingly, all three additional loci again exhibit very low rates of perfect repair (Supplementary Table 4). This indicates that the minor role of perfect repair may be a general feature of Cas9-induced DSBs, rather than a locus-specific phenomenon.

### Tight binding of Cas9 after cutting may explain erroneous and slow repair

We sought an explanation for the remarkably slow and error-prone repair that we observed. It was reported that the Cas9 can remain attached to the broken DNA ends after cutting *in vitro* (Sternberg et al., 2014), but it is not known how general this behaviour is. We therefore tested this for three DNA loci for which we had determined the kinetic rate constants. By PCR we first produced double-stranded DNA fragments of 600 – 1000 bp consisting of precisely the same sequences as the three loci. We then incubated each fragment *in vitro* with the respective Cas9/sgRNA complex to induce DSBs, and investigated the reaction products by agarose electrophoresis (Figure 6J, K).

The PCR product treated with Cas9/sgRNA-LBR2 showed the expected digestion products, but the smallest fragment was underrepresented while a broad smear indicated aberrant migration of the DNA. Strikingly, after heat denaturation of Cas9/sgRNA the smear disappeared and the smaller digestion product was clearly visible again. For sgRNA-AAVS1 and sgRNA-chr11 even more pronounced effects were observed: without denaturation the digested DNA appeared largely unbroken and the bands were shifted upwards, but after heat treatment it became clear that most of the DNA was in fact correctly digested. Together, these results indicate that the Cas9/sgRNA complex remains bound to the DNA ends after cutting, even when incubated overnight (Figure 6K). In the case of sgRNA-LBR2 it appears that this binding occurs primarily at one DSB end, while for the other two sgRNAs Cas9 remains bound to both ends. We propose that post-cutting adherence of Cas9 to the DNA ends impairs the repair process, thereby accounting for the slow and erroneous repair.

### Repair fidelity after double cutting

Finally, we investigated the fidelity of DSB repair upon induction of Cas9 in combination with two cotransfected sgRNAs that target adjacent sequences. If the two cuts are made simultaneously then the intermediate fragment may be lost, after which the two remaining ends are joined by the DSB repair machinery. The resulting junctions are amplified by PCR and sequenced in order to determine the error rate of the repair process, i.e., the frequency at which indels occur at the junction. Such a double cut strategy has been used previously in combination with I-SceI (Guirouilh-Barbat et al., 2007; Mao et al., 2008). Importantly, once the two ends are joined perfectly, they cannot be cut again because the new junction is not recognized by either of the two sgRNAs. This assay therefore complements our kinetic modeling of single-cut repair, in which cycles of repeated cutting and perfect repair were theoretically possible. We applied this double-cut strategy to further investigate the balance between perfect and imperfect repair.

We designed five sgRNAs targeting a second DSB site ~110-300bp upstream or downstream of the sgRNA-LBR2 target site. We then co-transfected each sgRNA together with sgRNA-LBR2, induced Cas9 expression and harvested cells after 60 hours. PCR amplification followed by high-throughput sequencing uncovered all possible intermediates and end-products that could be expected, such as junctions resulting from excision events as well as indels at one or both of the two cutting sites (Figure 7A–B).

**Figure 7:**
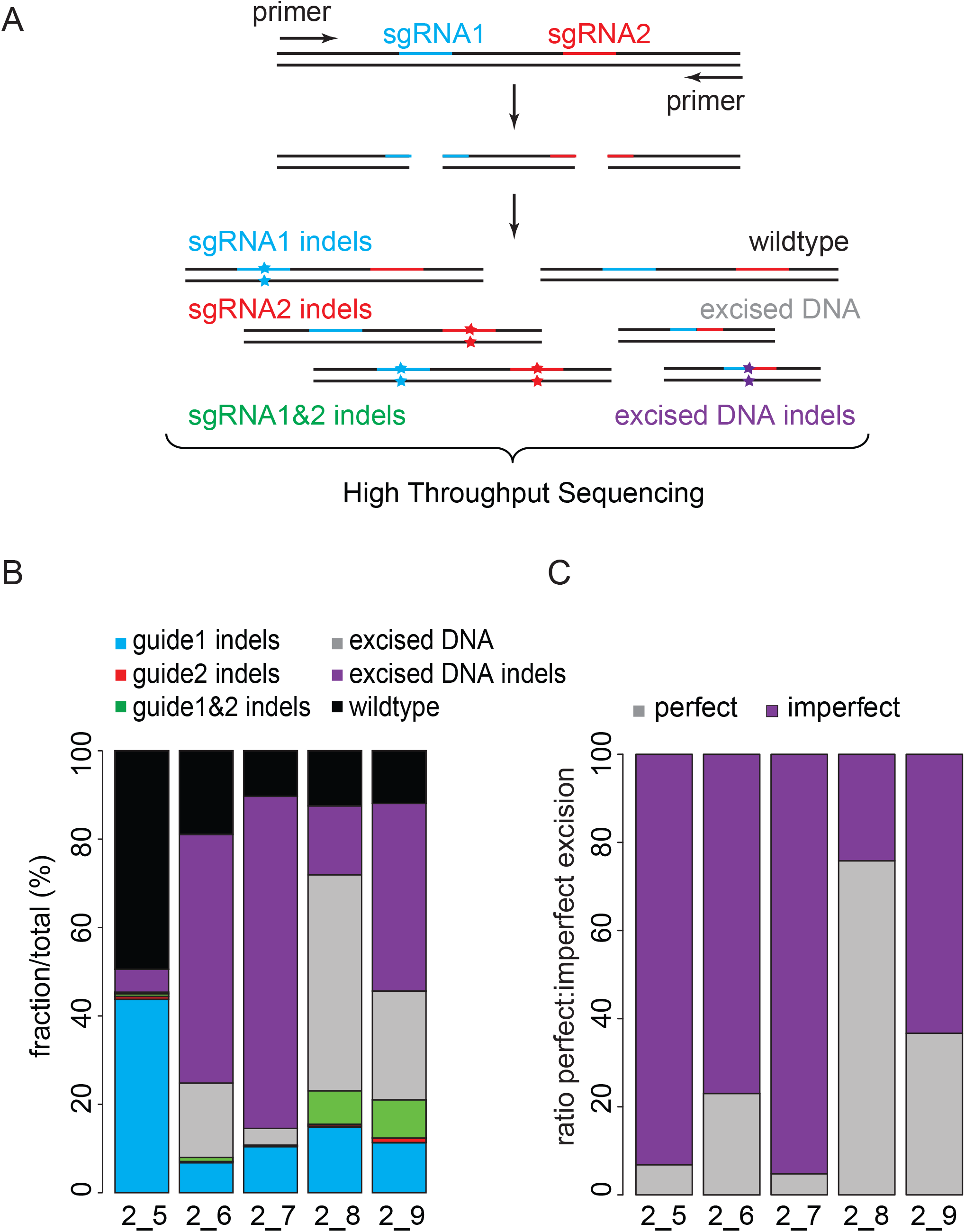
DSB repair fidelity after double cutting. (**A**) Repair products that may result from Cas9 in combination with two sgRNAs that target adjacent sequences. All products are amplified by PCR and detected by high-throughput sequencing. (**B**) Quantification of the various repair products 60 h after Cas9 induction in the presence of sgRNA-LBR2 and one of 5 sgRNAs (labeled 5 thru 9) that target a sequence within 110-300 bp from the LBR2 site. (**C**) Same as (**B**), but highlighting the relative proportions of perfect and imperfect junctions among the repaired excision events.

In those DNA molecules in which excision was successfully followed by repair, re-joining had occurred with highly varying degrees of fidelity. Depending on the combination of sgRNAs tested, we found that 25-95% of these junctions were imperfect (Figure 7C), as indicated by the occurrence of indels that ranged from 1 to ~20 bp in size. The variability in repair fidelity in this double-cut assay is much higher than we observed in the single-cut kinetic modeling, suggesting a complex, sequence-dependent course of events when two adjacent breaks occur.

Finally, in each tested double-cut combination we detected at least 2-fold more excised fragments than indels at single break sites (Figure 7B). This implies that the second cut typically occurred before the first cut had been repaired. This is in agreement with our modeling results indicating that repair of Cas9-induced breaks is a relatively slow process.

## Discussion

### DNA repair kinetics

CRISPR-Cas9 has rapidly become a widely used method for genome engineering. The precision, efficiency, and flexibility of this nuclease have been studied in detail to understand how it generates DSBs (Sander and Joung, 2014). However, the kinetics of re-joining of broken DNA ends after a Cas9-induced lesion has remained unclear. We therefore developed a strategy to directly measure the kinetics at single loci in human cells by direct quantitative measurements of intact and repaired DNA. Mathematical modeling of these data reveals that DSB repair of a Cas9-induced DSB is relatively slow (typically taking several hours) and has high error rates.

We found repair half-lives in the range of 5.0 to 8.7 h, which is substantially slower than previous estimates, which ranged from 0.25 to 2 h. These previous estimates were based on pulsed field gel electrophoresis, comet assays and tracking of repair foci after ionizing irradiation (Kinner et al., 2008; Metzger and Iliakis, 1991; Redon et al., 2009). These assays measured the disappearance of breaks and did not analyze the repair products. With our sequencing approach we can detect the repair products accumulating over time. In addition, most of these studies were performed after a pulse of ionizing radiation ranging from 1 – 20 Gy, which creates dozens to hundreds of DSBs. Here, we introduce at most 4 DSBs per cell (one target in each allele). We cannot rule out that a larger number of simultaneous DSBs accelerate the repair process. Although, this seems not very likely because in presence of 10 Gy IR damage in addition to the Cas9-induced DSB, we see only a small shift in repair pathway choice towards the faster C-NHEJ pathway.

Another factor that could contribute to the difference in repair rates is the possibility that Cas9 remains associated with the DNA ends after cleavage in cells, which could prevent access by the repair machinery. Our *in vitro* data show that Cas9 remains tightly bound to one or both DNA ends after cutting and rapid detachment could only be achieved by protein denaturation. This is in agreement with previous *in vitro* studies (Richardson et al., 2016b; Sternberg et al., 2014). *In vivo,* catalytically inactive Cas9 was also found to tightly bind to its target DNA (Knight et al., 2015), with a dwell time of about 2 hours (Ma et al., 2016, 2016). The bound Cas9/DNA complex may prevent recognition and binding of the DNA ends by the repair machinery.

### DNA repair fidelity

Our parameter estimates indicate that the majority of repair events after Cas9 cutting result in indels. This is surprising because C-NHEJ, which is typically the predominant pathway (Chiruvella et al., 2013; Wang et al., 2005), is generally thought to have low error rates (Betermier et al., 2014). The latter view was largely based on observations that error rates increase substantially in C-NHEJ-deficient cells compared to wild-type cells (Guirouilh-Barbat et al., 2004; Guirouilh-Barbat et al., 2007; Rass et al., 2009). We propose that the lingering Cas9 prevents error-free repair by C-NHEJ. This indirect mutagenic effect may be the key to the success of Cas9 as a genome editing tool.

The structure of the DNA ends could also affect the degree of accurate repair. High precision of C-NHEJ was found when DNA breaks were introduced by the I-SceI nuclease (Guirouilh-Barbat et al., 2004; Guirouilh-Barbat et al., 2007). I-SceI makes a staggered cut and leaves a 3’ overhang, while Cas9 generates blunt ends (Cong et al., 2013; Jinek et al., 2013; Mali et al., 2013). Blunt ends were shown *in vitro* and in wild-type yeast to be preferentially joined imprecisely (Boulton and Jackson, 1996; Burton et al., 2007; Daley and Wilson, 2005; Schar et al., 1997). Variants of Cas9 protein that generate DSBs with different overhangs (the nickases N863A and D10A) also resulted in differences in repair (Bothmer et al., 2017; Vriend et al., 2016). Thus, the high rates of imperfect repair that we observe may in part be related to the blunt ends created by Cas9.

The low rates of perfect repair also imply that HR plays only a minor role in the repair of Cas9-induced breaks. HR is known to be able to repair Cas9-induced breaks (Vriend et al., 2016) and K562 are proficient in HR (Voit et al., 2014). HR frequencies in combination with Cas9 have been estimated to be up to 4 – 15 % (Bothmer et al., 2017; Vriend et al., 2016), but these frequencies are likely to represent an overestimate because they were measured in the presence of a donor template that was either closely linked in cis or provided in excess by transient transfection. Normally HR only occurs in S and G2 phase of the cell cycle. We also cannot exclude that in those phases Cas9 is able to cut both sister chromatids at the same time and at the same site, thereby frustrating HR.

### No detectable rapid cycling

Our results do not support a rapid cycle of Cas9 cutting and perfect repair that terminates when a rare imperfect repair event occurs, as has been suggested (Richardson et al., 2016a). First, fitting of our model to the time series data is most consistent with slow and error-prone repair. Second, in a rapid cycle model the broken fraction would peak sooner than we observe. Third, rapid cycling is not compatible with the long residence time of Cas9 on its target DNA; *in vitro* Cas9 remains bound to the broken ends for several hours, while catalytically inactive Cas9 has a residence time of up to 2 hours *in vivo* (Ma et al., 2016, 2016; Richardson et al., 2016b, 2016b).

### Repair fidelity at double DSBs

The estimated ratios of perfect/imperfect repair rates were remarkably consistent between the four tested loci. In contrast, the proportion of perfect junctions in the double-cut assay was much more variable and strongly dependent on the precise combination of sgRNAs used. This suggests that repair of two nearby breaks is more complex than of a single break. For example, the first break may trigger local chromatin changes such as phosphorylation of H2A.X, which may in turn alter events at the second site, such as cutting rate and the recruitment of specific repair complexes. Furthermore, Cas9 may linger on one or both of the ends that are to be joined, and our *in vitro* data suggest that the dissociation rate depends on the sgRNA that is used. This may depend on the orientation of the PAM sites, or on other sequence features of the sgRNA used, but we have not been able to identify a predictive feature in the pairs of sgRNAs that we tested. Others have observed that sgRNA pairs resulted in high levels of precision repair (Canver et al., 2017; Geisinger et al., 2016; Zheng et al., 2014). We suggest that the presence of the bulky Cas9 protein at one or two ends can influence the accuracy of the joining of the two DNA ends.

### Different repair pathways

We found that at one genomic location both C-NHEJ and MMEJ can repair DSBs. We found that each of these repair pathways has different kinetics. The MMEJ operates with lower rates than C-NHEJ. This is in agreement with previous findings that indicate that C-NHEJ is the primary repair pathway (Chiruvella et al., 2013; Wang et al., 2005). This hypothesis is strengthened by the observation that MMEJ also exhibits a delayed onset compared to C-NHEJ. One possible interpretation is that under normal conditions the C-NHEJ system initially prevents access of the MMEJ pathway to the DSB; only after several hours, if C-NHEJ has failed to repair the break, the MMEJ pathway is allowed to engage. In contrast, upon inhibition of DNA-PKcs the MMEJ has immediate access and an increased rate of activity. These results are consistent with a previously proposed model in which MMEJ acts as a backup system for C-NHEJ (DiBiase et al., 2000; Guirouilh-Barbat et al., 2004; Guirouilh-Barbat et al., 2007; Wang et al., 2003; Wang et al., 2005).

### Effects of the local chromatin context

The local chromatin environment may affect Cas9 cutting efficiency as well as the repair outcome. Nucleosome position has an influence on the efficacy of Cas9 cutting and heterochromatin hampers the target search (Horlbeck et al., 2016; Isaac et al., 2016; Knight et al., 2015). Recruitment of repair factors is also affected by the chromatin environment (Aymard et al., 2014; Chiolo et al., 2013; Goodarzi and Jeggo, 2012; Goodarzi et al., 2008; Jakob et al., 2011; Janssen et al., 2016; Lemaitre et al., 2014; Tsouroula et al., 2016). We found that the cutting and repair rates are very similar across 4 different loci in the genome, of which three are located within active genes and one in an intergenic region. Certainly, we did not sample the full diversity of chromatin environments, and it will be interesting to test the repair kinetics and fidelity of many more loci in the genome to study how the rates are affected by chromatin environment.

## Author contributions

EKB designed the study, performed experiments, wrote code, analyzed data, wrote the manuscript. TC designed and conducted mathematical modeling, wrote code, analyzed data. MdH: optimized and performed experiments to measure DSBs; Cas9 western blots; *in vitro* Cas9 experiments. HAH contributed to studying the effects of IR and NU7441. WA wrote code; BvS designed and supervised the study, wrote the manuscript.

## Acknowledgments

We thank B. Evers for sharing plasmids prior to publication; H. te Riele and members of our laboratory for critical reading of the manuscript; the NKI Genomics and FACS core facilities for technical assistance. This work was supported by a ZonMW-TOP grant and ERC Advanced Grant 293662 (to BvS). WA was supported by an NWO-ALW grant awarded to M. van Lohuizen.

## Methods

### Cell culture and transfection

We established K562#17, which is a clonal cell line of K562 cells (American Type Culture Collection) stably expressing DD-Cas9. K562#17 cells were cultured in RPMI 1640 (Life Technologies) supplemented with 10% fetal bovine serum (FBS, HyClone®), 1% penicillin/streptomycin.

For transient transfection, 6x10^6^ K562 cells were resuspended in self-made transfection buffer (100 mM KH_2_PO_4_, 15 mM NaHCO_3_, 12 mM MgCl_2_, 8 mM ATP, 2 mM glucose (pH 7.4)) (Hendel et al., 2014). After addition of 3.0 μg plasmid DNA, the cells were electroporated in an Amaxa 2D Nucleofector using program T-016. DD-Cas9 was induced with a final concentration of 500 nM Shield-1 (Aobious).

For kinetics experiments, 18x10^6^ cells were transfected and divided over 12-well plates, one well for each time point and each well carrying 1x10^6^ cells. Cas9 was activated 24h after nucleofection and cells were collected at the indicated time points after Cas9 induction. As controls, cells without Shield-1 were collected at various time points.

DNA-PKcs inhibitor NU7441 (Cayman) final concentration of 1 μM or DMSO (control) was added to K562#17 at the same time when the cells were supplemented with Shield-1 to induce DD-Cas9. 10 Gy of IR was administered by Cs source Gammacell®40 Exactor (Best Theratronics).

### Cell viability

Cell viability was measured using a CellTiter-Blue Cell Viability Assay (Promega). CellTiter-Blue Reagent was 1:5 diluted in RPMI 1640 (Life Technologies) supplemented with 10% fetal bovine serum, 20 μL of this diluted CellTiter-Blue Reagent was added to 100 μL cell suspension. After a 3 hour incubation at 37 °C in a 96 well tissue culture plate, fluorescence (560_Ex_/590_Em_) was measured on an EnVision Multilabel Plate Reader (Perkin Elmer). Results of one CellTiter-Blue Cell Viability Assay per experiment are shown. The mean of two technical replicates is shown ± standard deviation (SD). Cell viability was measured 45 hours after Shield-1.

### Constructs

The sgRNA oligos (Supplementary Table 1) were cloned into expression vector pBluescript with the sgRNA cassette of PX330 (Addgene plasmid 42230) and transfected into K562#17. The sgRNAs were designed using CHOPCHOP (Montague et al., 2014). The pLenti-Cas9-T2A-Neo expression vector (Prahallad et al., 2015, 2015) was a kind gift of Dr. B. Evers, NKI. In the expression vector, the ubiquitin promoter was exchanged for the hPGK promoter and a destabilization domain (DD) (Banaszynski et al., 2006) was added at the N-terminus of the Cas9 gene, to generate DD-Cas9.

### High Throughput Sequencing

Cells were collected by centrifugation (300xg, 5 min) and the genomic DNA was isolated using the Isolate II Genomic DNA Kit (Bioline). PCR was performed in two steps; PCR1 with ~100 ng genomic DNA and site specific barcoded primers (see Supplementary Table 2). PCR2 used 2 μL of each PCR1 product with Illumina PCR Index Primers Sequences 1-12. Each sample was generated with a unique combination of a barcode and index. Both PCR reactions were carried out with 25 μL MyTaq Red mix (Bioline), 4 μM of each primer and 50 μL final volume in a 96 well plate. PCR conditions were 1 min at 95 °C, followed by 15 sec at 95 ° C, 15 sec at 58 ° C and 1 min at 72 ° C (15x). 20 μL of 8 samples were pooled and 100 μL was loaded onto a 1% agarose gel. PCR product was cut from gel to remove the primer dimers and cleaned with PCR Isolate II PCR and Gel Kit (Bioline). The isolated samples were sequenced by Illumina MiSeq. In this study, we only amplified the sgRNA on-target sequences. The effect of possible off-target activity of Cas9/sgRNA was ignored and is considered to be equal between the different experiments.

### LM-PCR

Genomic DNA (350 ng) was incubated with 0.1 μM primer EB479 and Phusion High Fidelity DNA polymerase (Thermo Fisher Scientific) for 5 min at 95 ° C, 30 sec at 55 °C and 30 sec at 72 ° C, to extend the non-blunt DNA ends near the break site. Subsequently, 0.16 μM dsDamID adaptor (Vogel et al., 2007) was ligated to the blunted broken ends at 16 ° C overnight using T4 ligase (5 U/μL, Roche). Ligase was heat inactivated for 20 minutes at 65 °C. To detect broken DNA a PCR was performed on the adaptor-broken DNA ligation product with 3 μM primer EB486 and EB487. In parallel an internal standard PCR was performed with the same samples using 3 μM primers EB488 and EB487 that are both located downstream of the sgRNA target site. PCR conditions were 4 min at 95 C, followed by 33 cycles of 10 sec at 95 ° C, 10 sec at 58 ° C, and 10 sec at 72 °C. During preparation the samples were always kept at 4 °C. The PCR products were analyzed on a 1.5% agarose gel and imaged with ChemiDoc™ Imaging Systems (BioRad). A box with a fixed volume was set over the appropriate LM-PCR band for each sample and the intensities within each box were determined in arbitrary units. These signals were divided by the intensities of the corresponding internal standard PCR signal. The signal at t = 0 h was considered background and was subtracted from all other time point samples. To correct for the arbitrary intensity values between different time series, each sample was divided by the sum of all values in one experiment. The mean of biological replicates is shown ± standard deviation (SD). The peak time (**τ**), when the broken fraction reaches a maximum, is determined by fitting the band intensity values of the time points with the expected curve shape from the three-state Ordinary Differential Equations (ODE) model.

### TIDE method

The TIDE method was performed as described in (Brinkman et al., 2014). Briefly, PCR reactions were carried out with ~100 ng genomic DNA in MyTaq Red mix (Bioline) and purified using the PCR Isolate II PCR and Gel Kit (Bioline). About 100-200 ng DNA from purified PCR samples was prepared for sequencing using BigDye terminator v3.1. Samples were analysed by an Applied Biosystems 3730x1 DNA Analyzer. The data obtained was analysed using the TIDE software (http://tide.nki.nl). The decomposition window used for TIDE was set to indels of size 0-10 bp.

### In vitro digestion with Cas9

PCR fragments of the target regions were amplified with myTaq (Bioline) according to manufacturer’s instructions. See Supplementary Table 2 for the used primers. *In vitro* transcribed sgRNA was generated by T7 promoter driven transcription using the Ribomax kit (Promega) and the RNA was purified with the MEGAclear kit (Life Technologies). 1.5 μL Cas9 (NEB), 50 μg sgRNA and 200 ng PCR fragment were incubated for 2 hours or overnight at 37 °C. For denaturation of the Cas9, samples were either incubated for 20 minutes at 80 °C or for 3 minutes at 96 °C and slowly cooled down to 20 °C (1 °C/min).

### Flow cytometry

K562#17 cells were collected 1 day after nucleofection and directly analyzed for fluorescence using a BD FACSCalibur. Viable cells were gated on size and shape using forward and side scatter.

For cell cycle profiles, cells were fixed with 5 mL of 70% ethanol at 4 °C overnight. After fixation, cells were washed with PBS and then incubated in PBS containing propidium iodide (PI) and RNase for 30 minutes at 37 °C in the dark. GFP & PI expression were measured using a 488 nm laser for excitation.

### Western blotting

Whole-cell extracts of ~0.5x10^6^ cells were prepared by washing cultures in PBS and lysing with 50 μL lysis buffer (Tris pH 7.6, 10% SDS, Roche proteinase inhibitor). Samples were pulse sonicated for 2 minutes and protein concentrations were determined using the Pierce BCA protein assay (Thermo Fisher Scientific). Samples containing 40 μg of total protein were separated by SDS-PAGE on 8% acrylamide gels and transferred to a nitrocellulose membrane through electroblotting. The membranes were blocked for 1 hour in PBS/Tween-20 0.1% containing 5% low fat milk. After washing twice with PBS/Tween20 0.1% the membranes were incubated with 1:2,000 α-Cas9 7A9 (Thermo Fisher Scientific) or γ-tubulin (T6558, Sigma) for 2 hours at room temperature with mild shaking. Subsequently, the membranes were washed again and incubated with a secondary antibody, 1:10,000 α-mouse IR800 (Li-Cor) at 37 °C for 1 hour with shaking. The antibody was detected by Odyssey scanner. The gel was analyzed by setting boxes with a fixed volume over the signals in each sample. The intensity within a set box was determined by Image Studio Version 2.0. Sample t = 60 h was loaded on every gel and used to normalize for the intensity of the signal in the different gels. The mean of three biological replicates is shown ± standard deviation (SD).

The relative activity of CRISPR-Cas9 is calculated by fitting the DD-Cas9 protein abundance from the quantified signals. The model employed for fitting describes stabilization of DD-Cas9 upon introduction of Shield-1, with the unit-less relative activity:

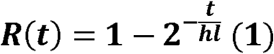

where *hl* is the protein half-life of DD-Cas9 with Shield-1 determined as 6.4 hours, and *t* is time after introduction of Shield-1. The model assumes that DD-Cas9 is very unstable without Shield-1, which is confirmed by the virtual absence of indel accumulation in the absence of stabilization (Figure 1E).

### NGS data analysis

In each sequence read, the distance between a fixed sequence at the start and at the end are determined and used to calculate a score, defined as the difference between the measured distance in the read and the expected distance in a wild-type sequence. *Insertions* and *deletions* have score >0 and <0, respectively. A *point mutation* has score=0, but some bases in the sgRNA target site differ from the wild-type sequence. The *intact* type specifies reads identical to wild-type sequences.

Per time point, the ratio of each type over the total of reads is calculated. We observed only 2 – 6.5% sequence reads in which we could not find a match with the constant parts and we discarded these reads in subsequent analyses. The called point mutations (score = 0) showed a very similar kinetic profile as the intact sequence (wild-type), indicating that they are mostly sequencing errors. We therefore assigned them as *intact* sequence in the analysis. *Insertion* and *deletion* levels at time point t = 0 h were considered as background and subtracted from all time points.

In the double-cut assays, paired end sequencing was performed. The forward and reverse read were matched by the unique sequence ID of a pair of reads. The deletion events were divided into two types: (i) perfectly excised DNA, and (ii) excised DNA with an indel when the deletion was larger than the expected excised product or up to 5 nucleotides smaller.

To determine the untransfected fraction, a standard sigmoid fit was applied to the time lapse curve of intact sequences according to the following equation.

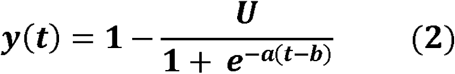

Where a, b and U are parameters that determine the shape of the curve. The fitting is done by the *nls* package of R by minimization of the Gaussian dispersion. U describes the asymptotic plateau, that is, the untransfected fraction.

### Mathematical modeling

#### I. Modeling the kinetics of total indels

The three fractions of intact (**P**), broken (**B**) and total indels (**M**) of a locus at any given time (t) after cutting by CRISPR-Cas9 must adhere to the principle of conservation

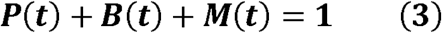

We assumed the cutting and repair kinetics is of first-order and the overall activity of CRISPR-Cas9 is proportional to the abundance of Cas9 that is modeled in equation (1). As shown in the diagram (Figure 1A) and taking equation (3) into account, the kinetics of the intact fraction (**P**) and the total indels (**M**) is determined by a nonlinear ODE

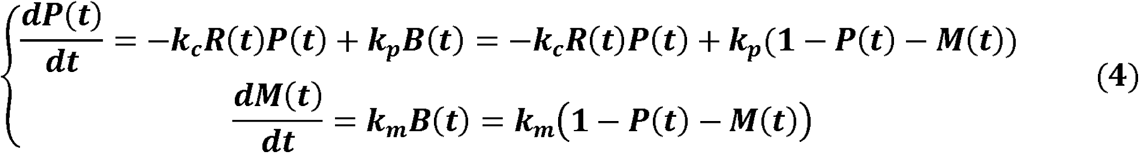

whereby *k_c_* stands for the maximal Cas9 cutting rate, *k_p_* for perfect repair rate and *k_m_* for mutagenic repair rate that gives rise to indels, all of which are in h^-1^.

Because broken DNA is not detected in amplicons across the break site, the measured intact fraction *(P_r_)* is the ratio between the abundance of intact sequences and the sum of the abundance of intact and indel sequences. Including the untransfected fraction (**U**) from equation (2) as a part in the intact fraction, the measured intact fraction is

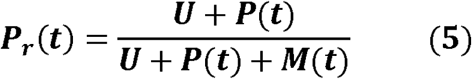

Taking equations (1), (2), (4) and (5) together, we estimated the kinetic parameters by minimizing the difference between the modeled P_r_(t) and experimentally measured data by the Levenberg-Marquardt algorithm (LMA) using Package FME in R. The kinetic profile of the broken fraction is then calculated according to equation (3).

#### II. Testing the robustness of the modelled perfect repair rate

For the parameter sweep analysis testing the robustness of the rate of perfect repair *k_p_* (Figure 2E, corresponding text), we restricted the ratio of *k_p_* to the total repair rate (*k_p_* + *k_m_*) as a sweeping factor

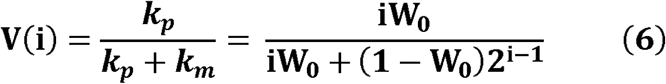

varying from W_0_ = 0.75 downwards by i times from 1 to 32, spanning 10 orders of magnitude. The incentive of introducing *i* as a term of multiplication is to test more carefully at high *k_p_* and a broad dynamic range with virtually equal step size at the low end. Therefore, we can represent *k_p_* as

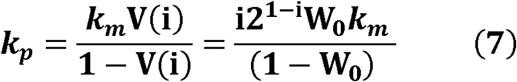

Taking equations (1), (2), (4), (5) and (7) together, we performed the model fitting with LMA.

Given a fixed ratio V(i) of perfect repair, the F-test was applied to examine the statistical significance for the difference in performance between restricted and optimal fittings, by the standard approach that takes the degree of freedom, which is the number of time points of the experiment minus 2 (the number of parameters in the model) and the fold difference in the deviance of fittings. A cutoff at p = 0.05 is applied to determine the upper bound of deviance and thereafter the upper bound of the ratio of perfect repair (V) is calculated accordingly (Figure 2E).

#### III. Modeling the kinetics of individual indels

To model +1 (*M*_+1_) and −7 (*M*_-7_) indels individually, we introduced the kinetic terms *k_+1_* and *k*_-7_ correspondingly for each mutant. As shown in the diagram (Figure 5A), the kinetics of individual indels can be written as

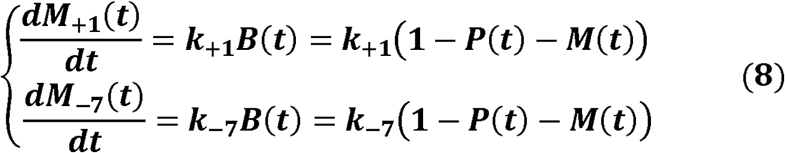

Similar to the measured intact fraction (*P_r_*),the measured fraction of individual indels (*M_r,+1_,M_r,-7_*) can be represented as

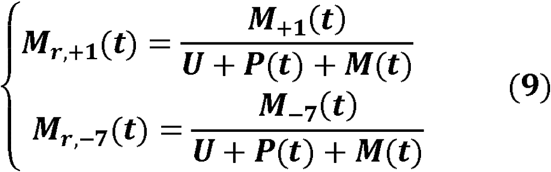

Taking equations (1), (2), (4), (5), (8) and (9) together, we estimated the kinetic parameters by minimizing a gradient of differences between the modeled P_r_(t), *M_r,_*_+1_(t) and *M_r,_*_-7_(t) and experimentally measured data by LMA.

#### IV. Adjustment of the modeling of the −7 indel kinetics

Following the modelling of individual indels, we discovered inconsistency between the fitted curve of the −7 indel and the experimental data (Figure 5C), suggesting the repair rate for the −7 indel is not a constant. Assuming a linear change of −7 repair rate over time, we adjusted the model by introducing a starting rate (*k*_-7,*t*=0*h*_) and an end rate (*k*_-7,*t*=60*h*_) (Figure 5D), and the non-constant −7 repair rate can be described as

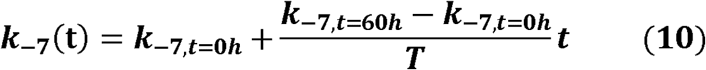

whereby *T* is 60 hours, the duration of experiment.

Adjusting equation (8) by equation (10), we have

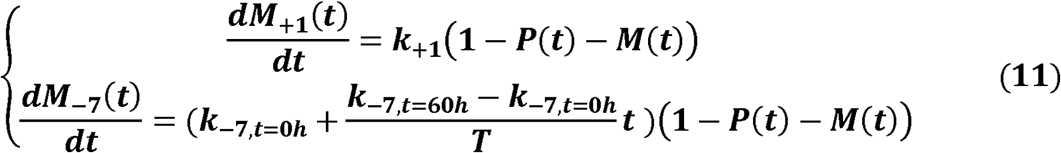

Taking equations (1), (2), (4), (5), (9) and (11) together, we performed the model fitting with LMA.

